# Regulation of temporal cytokine production by co-stimulation receptors in TCR-T cells is lost in CAR-T cells

**DOI:** 10.1101/2024.02.21.581341

**Authors:** Ashna Patel, Mikhail A. Kutuzov, Michael L. Dustin, P. Anton van der Merwe, Omer Dushek

## Abstract

CD8+ T cells contribute to immune responses by producing cytokines when their T cell receptors (TCRs) recognise peptide antigens on major-histocompability-complex (pMHC) class I. However, excessive cytokine production can be harmful. For example, cytokine release syndrome (CRS) is a common toxicity observed in treatments that activate T cells, including chimeric antigen receptor (CAR)-T cell therapy. While engagement of costimulatory receptors is well known to enhance cytokine production, we have limited knowledge of their ability to regulate the kinetics of cytokine production by CAR-T cells. Here we compare early (0-12 hours) and late (12-20 hours) production of IFN-γ, IL-2, and TNF-α production by T cells stimulated via TCR or CARs in the presence or absence ligands for CD2, LFA-1, CD28, CD27, and 4-1BB. For T cells expressing TCRs and 1st-generation CARs, activation by antigen alone was sufficient to stimulate early cytokine production, while co-stimulation by CD2 and 4-1BB was required to maintain late cytokine production. In contrast, T cells expressing 2nd-generation CARs, which have intrinsic costimulatory signalling motifs, produce high levels of cytokines in both early and late periods in the absence of costimulatory receptor ligands. Losing the requirement for costimulation for sustained cytokine production may contribute to the effectiveness and/or toxicity of 2nd-generation CAR-T cell therapy.

**Graphical abstract:** 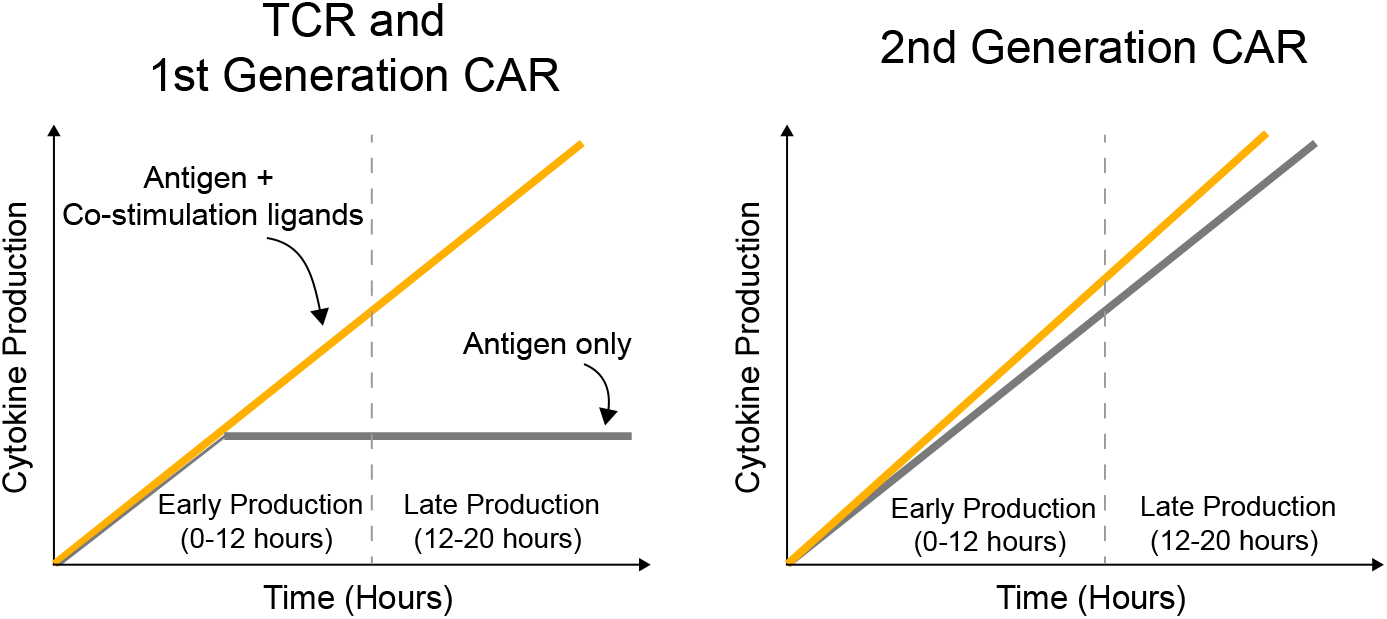

## Introduction

CD8^+^ T cells play central roles in immunity to intracellular pathogens and cancer. They can become activated to respond when their cell surface T cell receptors (TCRs) recognise peptide antigens displayed on major-histocompatibility-complex (pMHC) class I gene products on the surface of nearly all cells. The binding of pMHC to the TCR initiates a signalling cascade that can lead to multiple functional responses. These include the production of cytokines such as IFN-*γ*, IL-2, and TNF-*α* that collectively contribute to the ability of CD8^+^ T cells to execute immune responses (1, 2).

Because excessive cytokines levels can damage tissues, their production needs to be tightly regulated. One mechanism of regulation is engagement of co-stimulatory and/or co-inhibitory receptors on the T cell surface, which increases and decreases the production of cytokines, respectively (3–6). For example, cytokine production by T cells in cancer and chronic infections requires engagement of CD28 (7). We have limited information on the relative abilities of other co-stimulatory receptors, such as the adhesion receptors CD2 and LFA-1 and the TNFSF receptors CD27 and 4-1BB, to sustain T cell cytokine production.

The use of T cells in cell therapies to treat cancers, autoimmunity, and intractable chronic infections is showing great promise (8–10). In these therapies, T cells are transduced to express a new TCR (TCR-T) or chimeric antigen receptor (CAR-T) that recognises an appropriate target antigen before being infused back into patients. While TCR-T and 1st-generation CAR-T lack intrinsic co-stimulatory activity, 2nd-generation CAR-T contain co-stimulatory signalling motifs in their cytoplasmic domains. The role of costimulatory receptors and intrinsic co-stimulatory signalling motifs in CAR-T cell activation has previously been investigated (11–17) but their role in regulating sustained cytokine production by TCR-T and CAR-T cell remains poorly characterised. In the case of CAR-T cells, this is particularly important because patients often experience cytokine release syndrome (CRS), which can lead to organ failure and death (18).

We have previously developed and validated a reductionist system using TCR/CAR and co-stimulatory receptor ligands, and used it to investigate antigen sensitivity of TCRs and CARs (15, 17, 19–22) (Fig. 1A). In this work, we use the same system to investigate production of the cytokines IFN-*γ*, IL-2, and TNF- *α* (Fig. 1B). We found that, while cytokine production by TCR-T and 1st generation CAR-T cells was dependent on engagement of co-stimulation receptors, cytokine production by 2nd generation CAR-T cells was largely independent of co-stimulation receptors.

**Figure 1:**
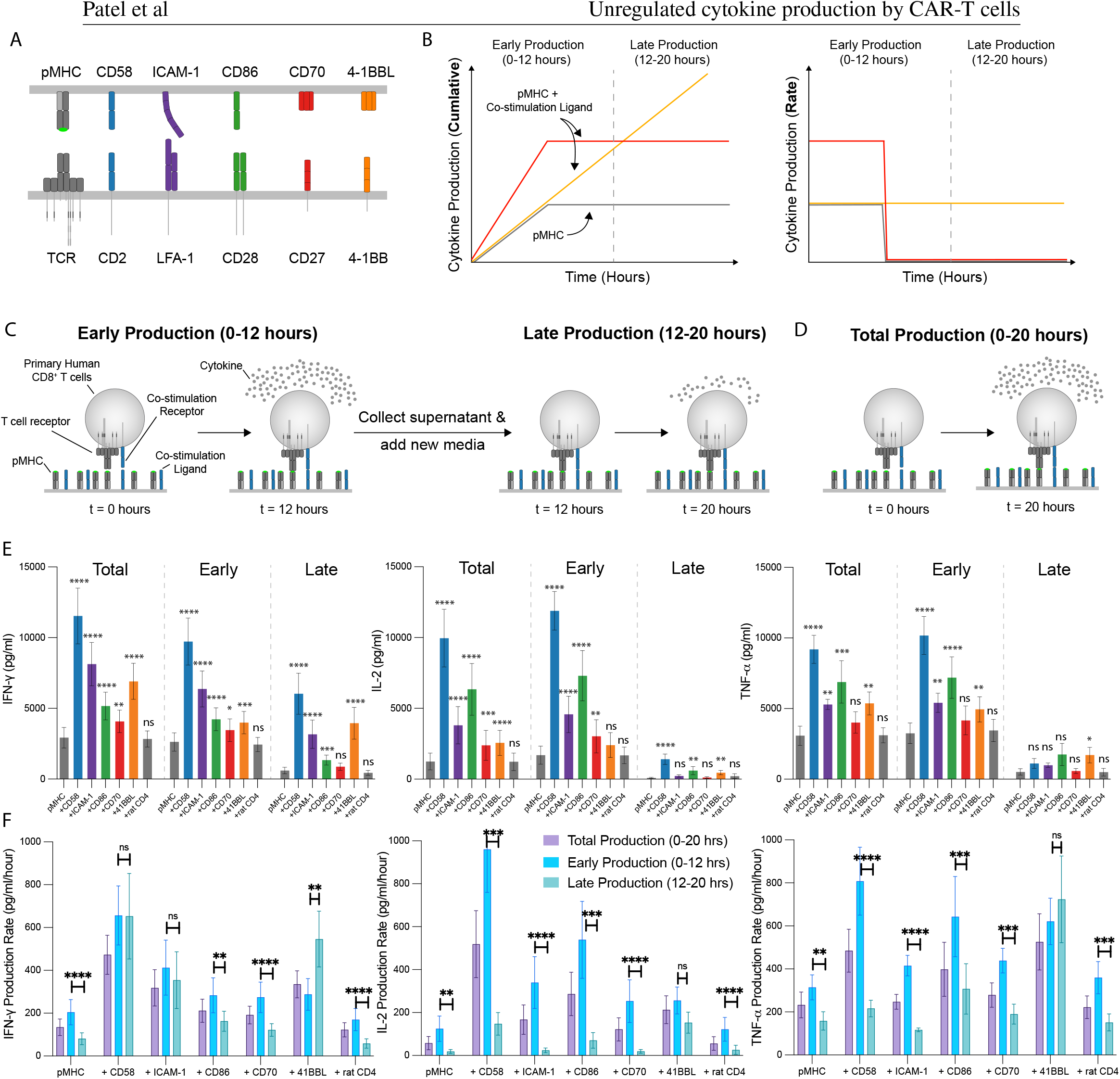
Regulated production of cytokine by co-stimulation receptors for the TCR. **(A)** Schematic of costimulation receptors and their ligands. **(B)** Schematics of potential cytokine production kinetics when T cells recognise antigen alone or with ligands to co-stimulation receptors. **(C**,**D)** Schematic of experiments to determine early, late, and total cytokine production. **(E)** Primary human CD8^+^ T cells transduced with the 1G4 TCR were stimulated by streptavidin surfaces coated with biotinylated pMHC alone or with the indicated biotinylated ligand. The supernatant level of the indicated cytokine was determined after 20 hours of stimulation (total production), after 12 hours (early production), or between 12 and 20 hours (late production). The rat CD4 ligand is used as a non-binding ligand control. Statistical significance of each ligand is compared to pMHC alone condition. **(F)** The production rate of each cytokine is calculated by dividing the supernatant cytokine data from (E) by the stimulation duration: 20, 12, or 8 hours for total, early, or late, respectively. Statistical significance was determined by multiple t-tests on log-transformed data with Holm-Sidak correction for multiple comparisons. Abbreviations: * = p-value⩽0.05, ** = p-value⩽0.01, *** = p-value⩽0.005, **** = p-value⩽0.001, ns = not significant. Data is shown as a mean +/-SEM of 6 independent experiments (i.e. 6 independent human donors).

## Results

### The temporal production of cytokines is tuned differently by different T cell co-stimulation receptors

To study the impact of T cell co-stimulation on the temporal kinetics of cytokine production, primary human CD8^+^ T cells transduced with the 1G4 TCR were stimulated with the cognate NY-ESO-1 pMHC antigen alone or together with the purified extracellular domain of ligands to the co-stimulation receptors CD2 (CD58), LFA-1 (ICAM-1), CD28 (CD86), CD27 (CD70), and 4-1BB (4-1BBL) (Fig. 1A). We coupled biotinylated antigen and ligands to streptavidin surfaces because this stimulation platform allowed for precisely defined surfaces that could persistently activate T cells without other compensatory/redundant molecules. We have previously observed (15, 21) that co-stimulation can prevent T cell desensitisation to TCR signals over a period of 20 hours leading to different outcomes for cytokine accumulation and production rates (Fig. 1B). In order to parse cytokine production kinetics the amount of IFN-*γ*, IL-2, and TNF-*α* was measured after 12 hours of stimulation (Early production), the media was then replaced and the amount of the same cytokines were measured over the next 8 hours (Late production) (Fig. 1C). As a control, we also measured the total cytokine produced after 20 hours without replacing the media (Total production)(Fig. 1D).

Consistent with their known costimulatory role, all tested costimulatory ligands increased total production of almost all cytokines (Fig. 1E). The single exception was that CD70 did not increase TNF-*α* production. The CD2 costimulatory ligand CD58 had the largest impact on the production of all 3 cytokines. The second most potent costimulatory ligand varied with cytokine: it was the LFA-1 ligand ICAM-1 for IFN-*γ* and the CD28 ligand CD86 for IL-2 and TNF-*α*.

Interestingly, distinct effects were seen when comparing early versus late production (Fig. 1E), especially for IL-2 and TNF-*α*. While most ligands increased early production of each cytokine, only some increased late production. For example, CD2, LFA-1, CD28, and 4-1BB engagement increased early production of TNF-*α* but only 4-1BB engagement increased late production.

To directly compare early and late production, we calculated the cytokine production rate by dividing by the collection duration (Fig. 1F). The rate of cytokine production decreased between early and late production for all three cytokines when T cells were stimulated with pMHC antigen alone. Interestingly, co-stimulatory ligands differed in their effect on late cytokine production. 4-1BB co-stimulation maintained the production rate of IL-2 and TNF-*α*, and increased the production rate of IFN-*γ*. CD2 and LFA-1 costimulation maintained the production rate of IFN-*γ*, but not IL-2 and TNF-*α*. Finally, CD28 and CD27 co-stimulation failed to maintain the production rate of any of the three cytokines.

Previous work has suggested that T cells can tune their cytokine production based on feedback from extracellular cytokine levels (23, 24). To exclude this we examine the effect of varying the density of T cells in the culture (Fig. S1). Our finding that cytokine levels scaled linearly with cell numbers shows that cytokine production is not affected by extracellular cytokine levels, ruling out feedback effects.

Taken together, these results indicate that sustained (20 h) cytokine production induced by TCR engagement requires co-stimulation, and that co-stimulatory receptors differ in their impact on cytokine production.

### Unregulated cytokine production by 2nd generation CAR-T cells

We next compared early and late production of cytokines by CAR-T cells. We used a previously described 1st generation CAR containing the *ζ*-chain cytoplasmic domain and a 2nd generation CARs contain both the *ζ*-chain and CD28 cytoplasmic domains (17). These CARs use the D52N scFv, which like the 1G4 TCR, recognise the NY-ESO-1 pMHC (Fig. 2A). The D52N scFv is derived from the 3M4EF Fab, which binds pMHC in the same orientation as the TCR (25, 26). Using pMHC tetramers, we confirmed that both CARs expressed at similar levels to the TCR (Fig. 2B).

**Figure 2:**
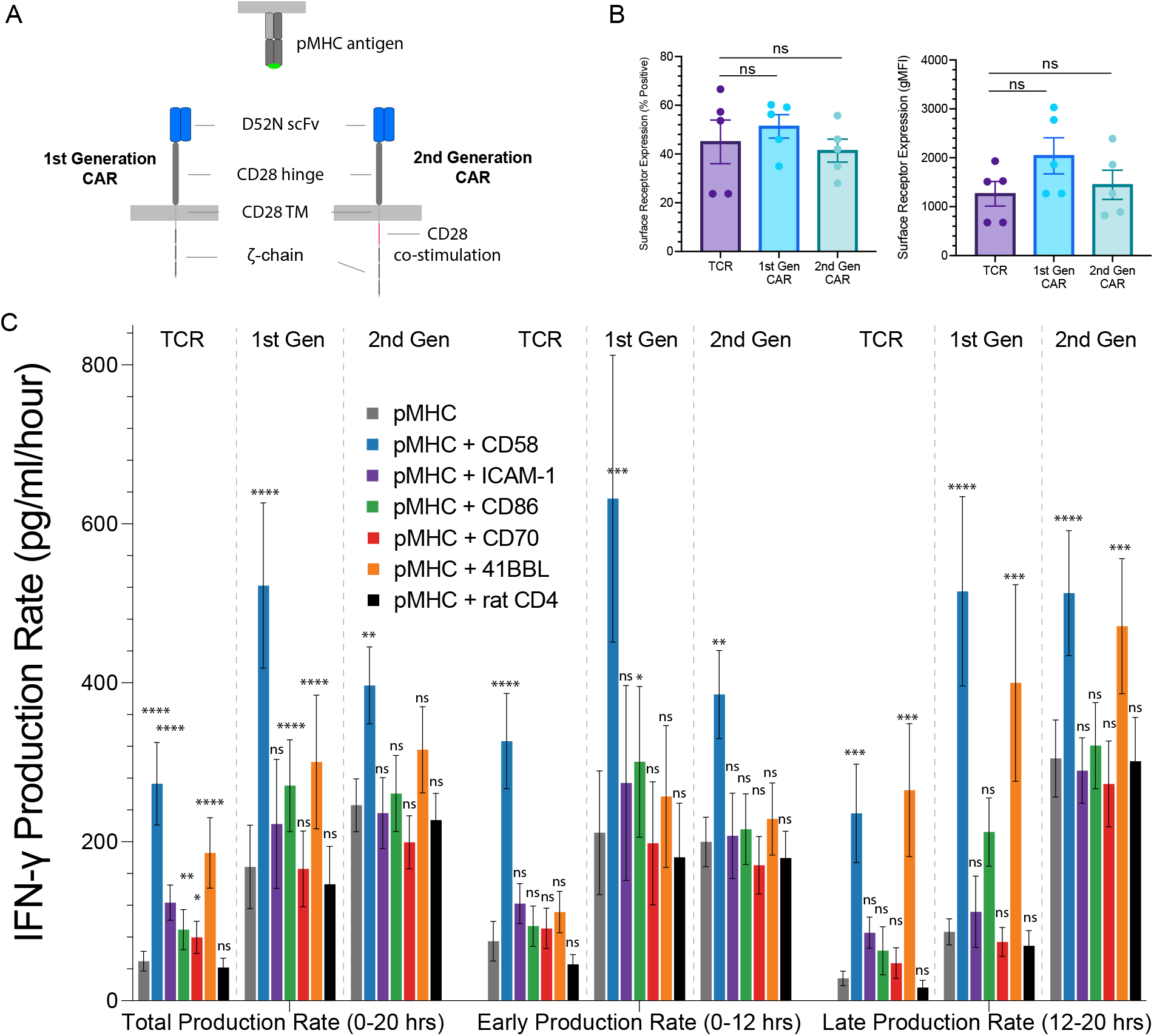
Regulated production of the cytokine IFN-*γ* by co-stimulation receptors for 1st generation CAR-T cells is reduced for 2nd generation CAR-T cells. **(A)** Schematic of 1st and 2nd generation CARs that use the D52N scFv to recognise the NY-ESO-1 pMHC. **(B)** Surface expression of each antigen receptor on transduced primary human CD8^+^ T cells is detected using pMHC tetramers in flow cytometry. **(C)** Rate of IFN-*γ* production for the indicated conditions. Statistical significance was determined by multiple t-test on log-transformed data relative to the pMHC alone condition with a Holm-Sidak multiple comparisons correction. Abbreviations: * = p-value⩽0.05, ** = p-value⩽0.01, *** = p-value⩽0.001, **** = p-value⩽0.0001, ns = not significant. Data is shown as a mean +/-SEM of 5 independent experiments (i.e. 5 independent human donors).

We measured total, early, and late cytokine production by TCR and CAR expressing CD8^+^ T cells in response to pMHC antigen alone or with co-stimulation ligands. CD2 and 4-1BB engagement had the largest impact on IFN-*γ* production, particularly for late production (Fig. 2C), while CD2, CD28 and 4-1BB engagement enhanced IL-2 and TFN-*α* production (Fig. 3).

**Figure 3:**
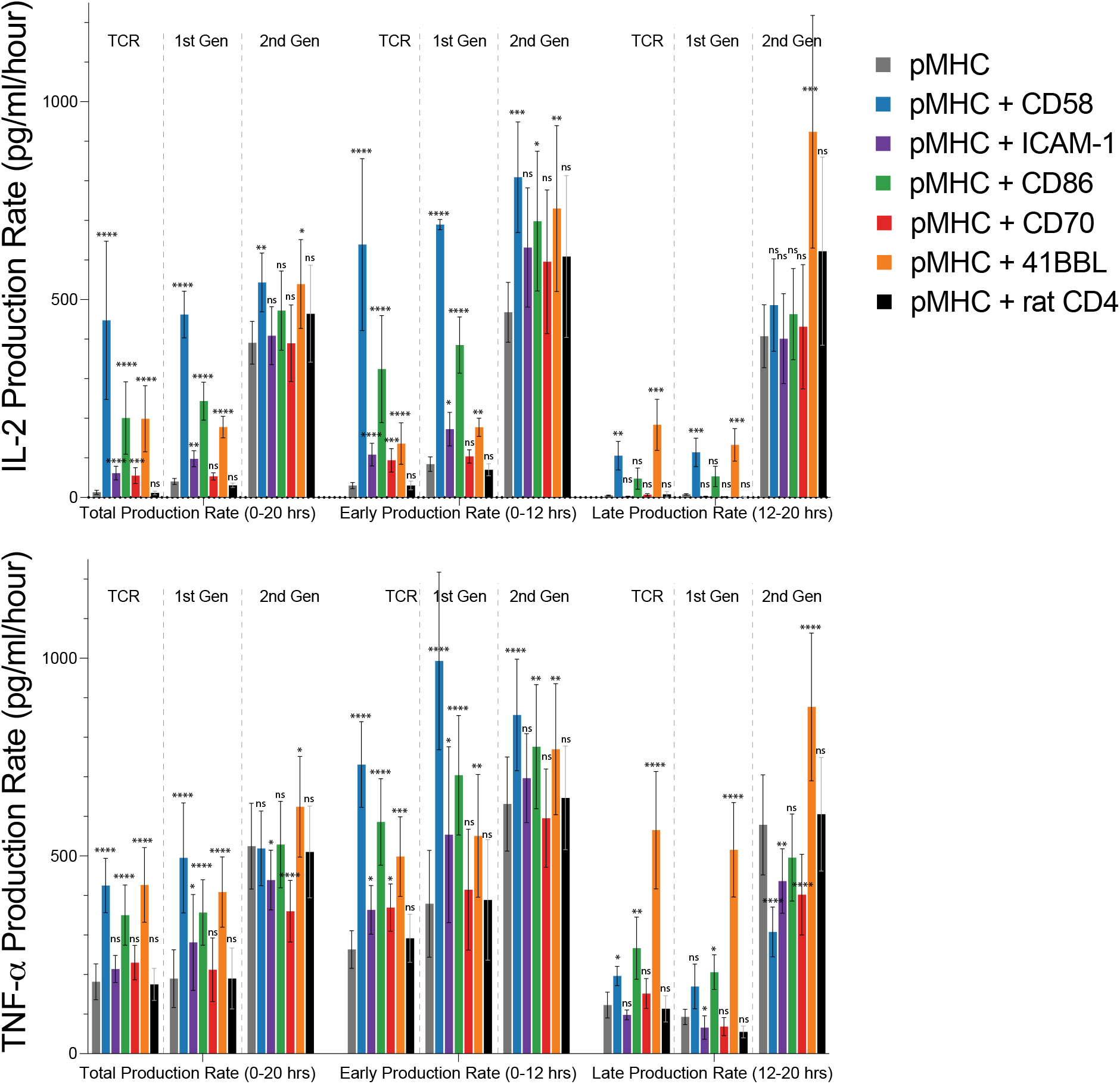
Regulated production of the cytokine IL-2 and TNF-*α* by co-stimulation receptors for 1st generation CAR-T cells is reduced for 2nd generation CAR-T cells. The production rate of IL-2 (top) and TNF-*α* (bottom) for the indicated antigen receptor, production period, and co-stimulation ligand. Statistical significance was determined by multiple t-test on log-transformed data relative to the pMHC alone condition with a Holm-Sidak multiple comparisons correction. Abbreviations: * = p-value⩽0.05, ** = p-value⩽0.01, *** = p-value⩽0.001, **** = p-value⩽0.0001, ns = not significant. Data is shown as a mean +/-SEM of 5 independent experiments (i.e. 5 independent human donors).

Interestingly T cells expressing a 2nd generation CAR had a striking ability to maintain late cytokine production rates in response to antigen alone (Figs. 2C and 3). This is more easily observed when plotting early cytokine production against late cytokine production (Fig. 4). In the case of the TCR and 1st generation CARs, the production rate of all cytokines decreased between early and late periods in response to antigen alone. Similarly, extrinsic co-stimulation by CD2 and particularly 4-1BB was required to maintain cytokine production. In striking contrast, 2nd generation CARs were able to maintain a high rate of cytokine production in response to antigen alone, even without co-stimulation. Moreover, LFA-1 and CD27 did not increase cytokine production by 2nd generation CARs and the modest increase induced by CD28 was only observed on early cytokine production.

**Figure 4:**
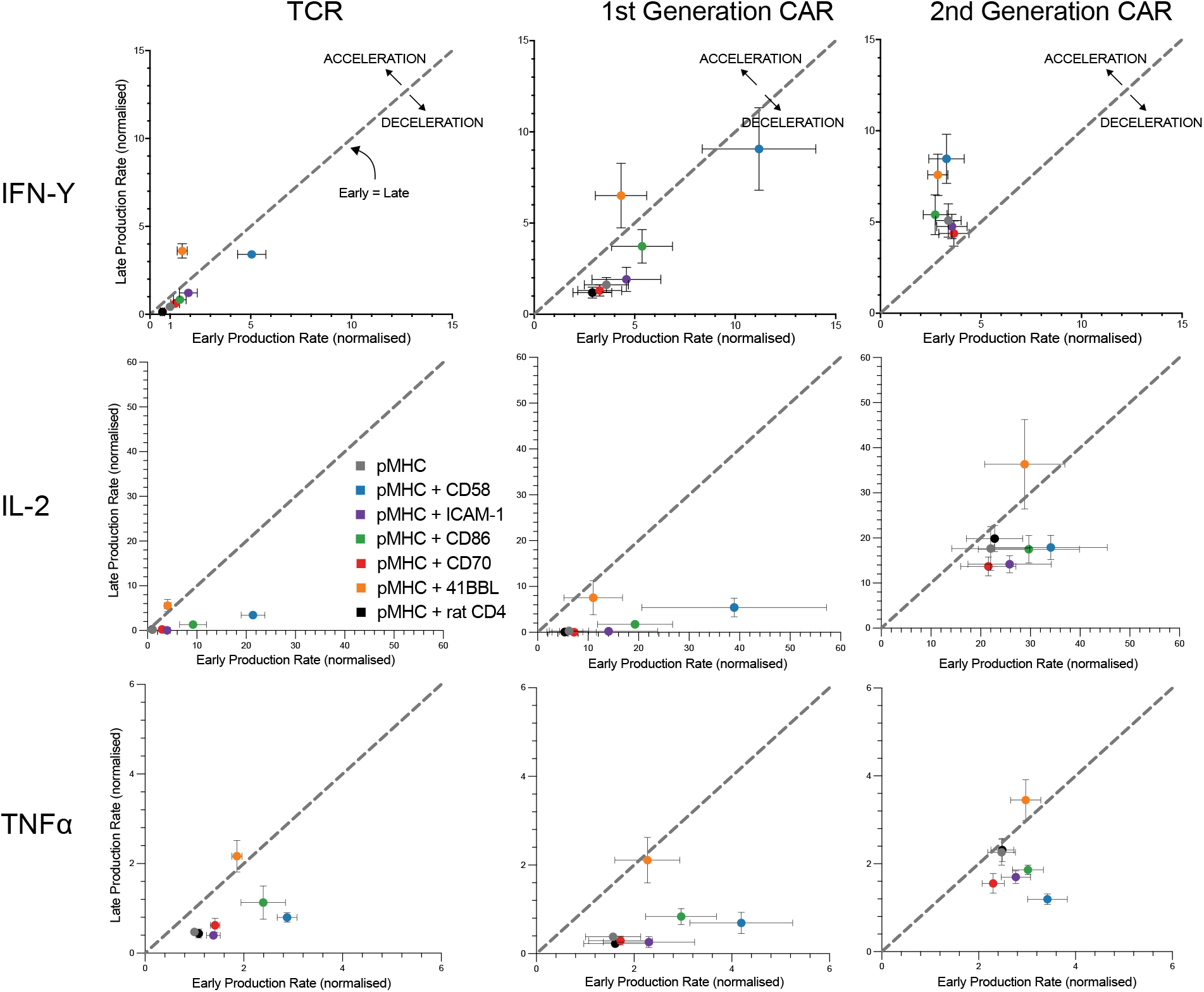
Cytokine production rates are maintained by 2nd generation CAR-T cells largely independent of extrinsic co-stimulation. The late production rate is plotted over the early production rate for the indicated cytokine (rows) and antigen receptor (columns). All data is replotted from Fig. 2-3 normalised to the early rate of cytokine production by the TCR in response to pMHC alone. All statistical analysis can be found in Fig. 2-3. Data is shown as a mean +/-SEM of 5 independent experiments (i.e. 5 independent human donors).

Taken together these results show that 2nd generation CARs stimulate more sustained cytokine production than 1st generation CARs and TCRs that is less dependent on costimulation.

## Discussion

Using a reductionist system, we have found that cytokine production by T cells is highly regulated by extrinsic co-stimulation ligands when recognising antigen through the TCR or 1st generation CAR. However, 2nd generation CARs displayed largely unregulated cytokine production that is maintained independent of extrinsic co-stimulation receptors.

The ability of T cells to orchestrate immune responses relies on finely tuned production of cytokines. This may be important to enable the clearance of pathogens without excessive tissue damage (7). We found that the rate of cytokine production decreases when T cells recognise antigen in isolation, as in our previous work (15, 21). As a result, T cells become increasingly reliant on signalling from co-stimulation receptors to sustain cytokine production. The sustained production of IFN-*γ*, IL-2, and TNF-*α* over 20 hours could be achieved by 4-1BB co-stimulation whereas co-stimulation by CD2 and LFA-1 could only sustain IFN-*γ* production. Although CD28 and CD27 co-stimulation increased the early rate of cytokine production (except TNF-*α* for CD27), they could not sustain this higher rate with CD27 co-stimulation completely absent over the late production period. The impact of all of these co-stimulation receptors have previously been shown to be TCR dependent (15, 21, 27–29), although in the case of 4-1BB there is also evidence for a TCR independent signalling function (30, 31), which may explain its unique ability to sustain cytokine production. We have used expanded primary human CD8^+^ T cells and a similar ‘wave’ of cytokine production has been observed in a variety of un-expanded primary T cells, including naive, central memory, effector memory, when stimulated by the TCR and CD28 (32). Taken together, these results suggest that sustained cytokine production by T cells relies on extrinsic co-stimulation receptors.

In contrast to the TCR, we found that antigen recognition by 2nd generation CARs produced largely unregulated cytokine production. Whereas the TCR and 1st generation CARs required extrinsic ligation of co-stimulation receptors to increase and sustain cytokine production, 2nd generation CARs maintained (IL-2, TNF-*α*) or increased (IFN-*γ*) their rate of cytokine production over time independent of extrinsic co-stimulation. The only detectable regulation was by 4-1BB which led to an even higher production rate of all cytokines, which may be a result of the non-overlapping signalling pathways induced by CD28 and 4-1BB (3, 30). The unregulated production of cytokines by CAR-T cells is a double edged sword. On the one hand, cancer cells are known to reduce expression of ligands to co-stimulation receptors to escape T cell immunity. As a result, reducing the dependence on co-stimulation ligands for cytokine production is a useful feature reducing the ability of cancers to evade CAR-T cell responses. On the other hand, the inability to extrinsically control CAR-T cells means that the normal regulatory processes of adaptive immunity are lost increasing the possibly of cytokine storms leading to excessive inflammatory responses and tissue damage.

## Acknowledgements

We thank Johannes Pettmann and John Nguyen for assistance in protein production.

## Funding

The work was funded by a Wellcome Trust Senior Fellowship in Basic Biomedical Sciences (207537/Z/17/Z to OD), by a Biotechnology and Biological Sciences Research Council (BBSRC) studentship (grant number BB/M011224/1 to AP), and by a Kennedy Trust for Rheumatology Research Professorship (to MLD).

### Open access

This research was funded in whole, or in part, by the Wellcome Trust [207537/Z/17/Z]. For the purpose of Open Access, the author has applied a CC BY public copyright licence to any Author Accepted Manuscript version arising from this submission.

## Data Availability

The data underlying this article are available in the article and in its online supplementary material.

## Author contributions

Conceptualization (OD), Data Curation (AP, OD), Formal Analysis (AP, OD), Funding Acquisition (OD), Investigation (AP, MAK, MLD, PAvDM, OD), Methodology (AP), Project Administration (OD), Supervision (MLD, PAvDM, OD), Visualization (AP, OD), Writing – Original Draft (AP, OD), Writing – Review & Editing (AP, MLD, PAvdDM, OD)

## Materials & Methods

### Protein Production & Purification

#### Ligands for co-stimulation receptors

The soluble extracellular domain (ECD) fused to a c-terminus His-tag and AviTag of human CD58, ICAM-1, CD86, 4-1BBL and CD70 were produced in adherent HEK 293T cells. Plasmids encoding TNFSR ligands (4-1BBL and CD70) were a kind gift from Harald Wajant (Würzburg, Germany) and incorporate a Flag tag and tenascin-C trimerisation domain. Adherent HEK 293T cells were transfected with Xtremegene HP reagent (Roche) according to manufacturer’s protocol. Supernatants were filtered using a 0.45*µ*M filter and proteins purified on a Ni-NTA agarose column. Biotinylation of proteins was performed either *in vitro* as described above, or *in situ* through co-transfection of BirA encoding plasmid (10%) with 100*µ*M D-biotin. Additional purification with removal of excess biotin was performed through size-exclusion chromatography in HBS-EP. Purified proteins were stored at -80°C.

#### Peptide-MHC Monomers

The 9V variant (SLLMWITQ**V**) derived from the wild-type NY-ESO-1_157*−*165_ 9C peptide was synthesised at a purity of *>* 95% (Peptide Protein Research, UK). Class 1 pMHCs were produced by expressing Human HLA-A*0201 heavy chain and human *β*-2microglobulin with a C-terminal BirA tag in *E*.*coli* as inclusion bodies and refolded with the 9V peptide *in vitro* as described previously (22, 33). Protein was biotinylated using the BirA enzyme (Avidity, USA) and purified via size-exclusion chromatography (Superdex S75 column, GE Healthcare, USA), in HBS-EP buffer (10 mM M HEPES pH 7.4, 150 mM NaCl, 3 mM EDTA, 0.005% v/v Tween-20). Purified protein was aliquoted and stored at -80°C until use.

#### Peptide-MHC Tetramers

Fluorescently conjugated pMHC tetramers were produced to enable detection of antigen receptors (TCR/CAR). Streptavidin-PE was gradually added to biotinylated 9V pMHC at a 4:1 Molar ratio, whilst shaking. Tetramers were stored for up to 3 months at 4°C in the dark.

### Production of TCR or CAR transduced primary human CD8+ T cells

#### Primary Human CD8+ T Cell Isolation

Human CD8+ T cells were isolated from leukocyte cones purchased from the National Health Service’s (UK) Blood and Transplantation service. Isolation was performed using negative selection. Briefly, blood samples were incubated with Rosette-Sep Human CD8+ enrichment cocktail (Stemcell) at 150 for 20 minutes. This was followed by a 3.1 fold dilution with PBS before layering on Ficoll Paque Plus (GE) at a 0.8:1.0 ficoll to sample ratio. Ficoll-Sample preparation was spun at 1200for 20 minutes at room temperature. Buffy coats were collected, washed and isolated cells counted. Cells were resuspended in complete RMPI (RPMI supplemented with 10% v/v FBS, 100 penicillin, 100 streptomycin) with 50U of IL-2 (PeproTech) and CD3/CD28 Human T-activator Dynabeads (Thermo Fisher) at a 1:1 bead to cell ratio. At all times isolated human CD8+ T cells were cultured at 37 and 5% CO2.

#### Lentivirus Production

HEK 293T cells were seeded in DMEM supplemented with 10% FBS and 1% penicilin/streptomycin in 6-well plates to reach 60–80% confluency on the following day. Cells were transfected with 0.25 pRSV-Rev (Addgene, 12253), 0.53 *µ*g pMDLg/pRRE (Addgene, 12251), 0.35 *µ*g pMD2.G (Addgene, 12259), and 0.8 µg of transfer plasmid using 5.8 X-tremeGENE HP (Roche). Media was replaced after 16 hours and supernatant harvested after a further 24 hours by filtering through a 0.45 cellulose acetate filter. Supernatant from one well of a 6-well plate was used to transduce 1 million T cells.

#### TCR & CAR Transduction

1 million cells in 1mL of complete RPMI were grown overnight in TC-treated 12 well plates. On the following day T Cells were transduced using lentivirus encoding for 1G4 TCR or CAR contructs. On days 2 and 4 post-transduction, 1 mL of media was exchanged and IL-2 was added to a final concentration of 50 Units/mL. Dynabeads were magnetically removed on day 5 post-transduction. T cells were then cultured at a density of 1 million cells/mL and supplemented with 50 Units/mL IL-2 every other day. T cells were used between 10 and 16 days after transduction.

#### Plate Stimulation Assay

Biotinylated pMHC and biotinylated costimulatory ligand (CD58, ICAM-1 or CD86) were diluted to the required concentration and added to Pierce Streptavidin Coated High Capacity 96 well plates (Thermo Fisher) in 50*µ*L PBS and incubated for 45 minutes at room temperature or left overnight at 4 °C.

The 96-well plates were the washed twice in PBS to remove excess protein. CD8^+^ T cell blasts T cells transduced with TCR or CAR, were counted and washed before being added to wells in the 96 well plate. 75,000 T cells were added in 200*µ*L complete RPMI unless otherwise stated. Cells were spun briefly at 50g, 37 °C for 1 minute to allow them to settle at the bottom of each well and make contact with relevant protein, before incubation for the required time at 37 °C 5% CO_2_. At the end of the stimulation assay, the supernatant was carefully removed and saved for ELISA analysis. 10mM EDTA in PBS was then added to detach the T cells. The cells were then aspirated and transferred to a v-bottom plate and washed once in 200*µ*L PBS 1% BSA (500g, 4°C, 5 minutes). Antibodies against T cell activation markers were diluted in PBS 1% BSA at a 1:200 dilution. To detect TCR-CAR expression fluorescently-conjugated peptide-MHC tetramers were added to the staining antibodies at a 1:1000 dilution. A viability dye was also added at a dilution if 1:2500 to distinguish live cells from dead cells. 50*µ*L of this staining solution was to the cells, before incubating them for 20 minutes at 4°C in the dark. The cells were washed twice in PBS, and resuspended in 75*µ*L PBS, before running on a flow cytometer. Flow cytometry data was analysed using FlowJo (BD Biosciences)

#### Cytokine detection

IL-2 Human uncoated ELISA kit, TNF-*α* Human uncoated ELISA kit, or IFN-*γ* Human uncoated ELISA kit and Nunc MaxiSorp 96-well plates were used according to the manufacturer’s instructions. The supernatant from stimulation assays was diluted 1 in 15 for ELISAs. The absorbance at 450 nm and 570nm were measured using a SpectraMax M5 plate reader (Molecular Devices).

#### Data Analysis

All statistical tests were performed in PRISM V.9 (GraphPad Software). ELISA standard curves were fitted using linear regression.

## Supplementary Information

**Figure S1:**
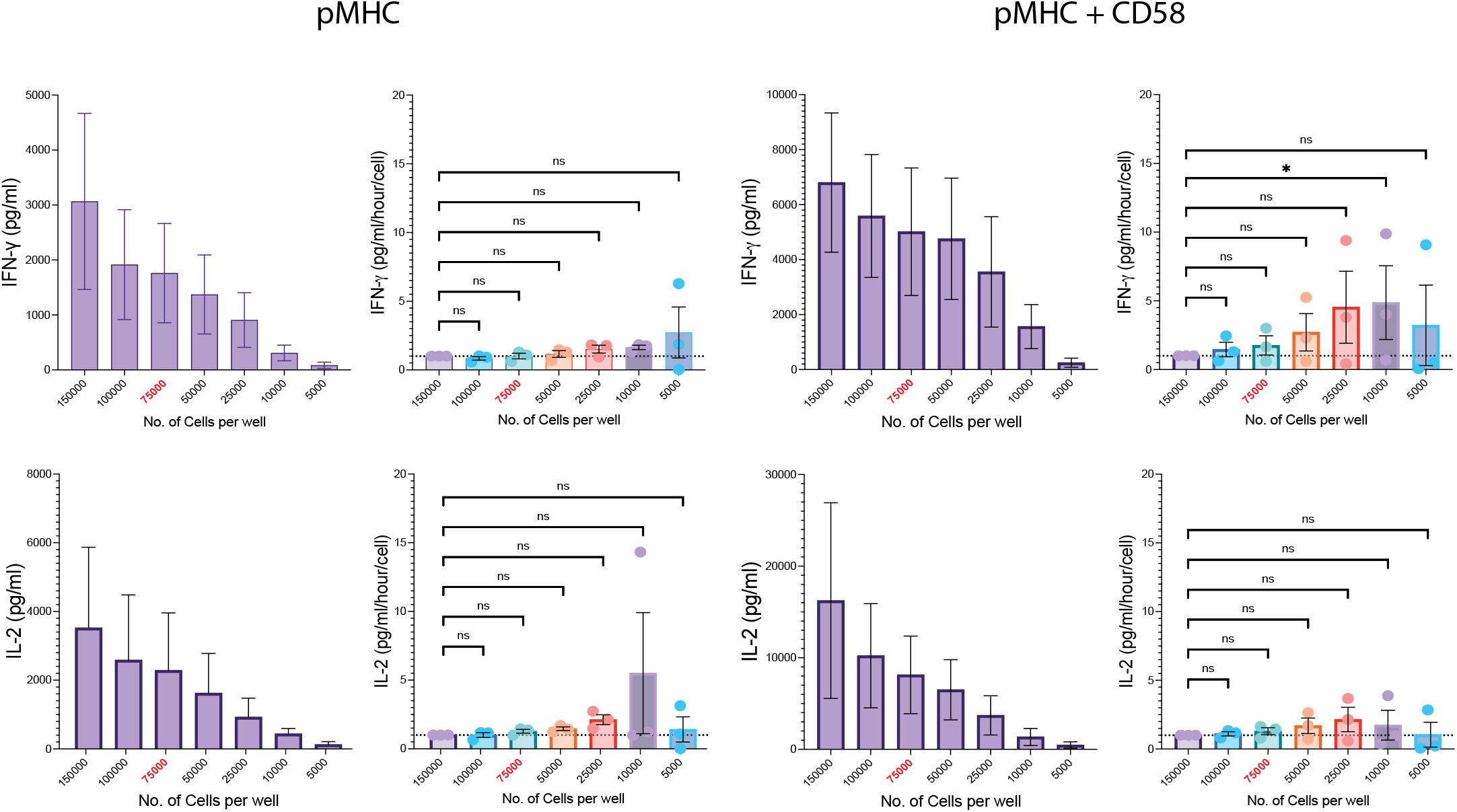
Cytokine production scales linearly with T cell number. The indicated number of primary human CD8^+^ T cells transduced with the 1G4 TCR (from 150,000 per well to 5000 per well) were stimulated by pMHC or by pMHC and CD58 for 20 hours before measuring IFN-*γ* (top row) or IL-2 (bottom row). All other experiments in the manuscript were performed with 75,000 cells per well (indicated in red). Statistical significance was determined by 1-way ANOVA on log-transformed data. Abbreviations: * = p-value⩽0.05.

